# Cell surface morphology mimicking nano-bio platform for immune cell stimulation

**DOI:** 10.1101/2024.01.23.576453

**Authors:** Beena Varghese, José Alfredo González-Navarro, Valentino Libero Pio Guerra, Margarita Faizulina, Daria Artemieva, Tomáš Chum, Tejaswini Rama Bangalore Ramakrishna, Marek Cebecauer, Petr Kovaříček

## Abstract

Studying the complex realm of cellular communication and interactions by fluorescence microscopy requires sample fixation on a transparent substrate. To activate T cells, which are pivotal in controlling the immune system, it is important to present the activating antigen in a spatial arrangement similar to the nature of the antigen-presenting cell, including the presence of ligands on microvilli. Similar arrangement is predicted for some other immune cells. In this work, immune cell-stimulating platform based on nanoparticle-ligand conjugates have been developed using a scalable, affordable, and broadly applicable technology, which can be readily deployed without the need for state-of-the-art nanofabrication instruments. The validation of surface biofunctionalization was performed by combination of fluorescence and atomic force microscopy techniques. We demonstrate that the targeted system serves as a biomimetic scaffold on which immune cells make primary contact with the microvilli-mimicking substrate and exhibit stimulus-specific activation.

## Introduction

T cells are an essential part of the immune system protecting vertebrates from infection or cancer and their misfunction may lead to diverse pathologies, e.g., immunodeficiency or autoimmunity diseases. T cells are activated and regulated through their characteristic surface receptors. T cell receptor (TCR) recognizes foreign patterns (antigens) in the form of the peptide-major histocompatibility complex (pMHC) on the surface of target cells called antigen presenting cells (APCs). Productive interaction triggers an initial activation of signalling.^1^ In almost all published T-cell activation studies employing advanced fluorescence microscopy, the APC was represented by the optical surface (coverslip) and ligands restricted to the focal plane by direct immobilisation on the glass or via a supported planar bilayer. In other words, T cells were always interacting with a flat surface in these experiments. Such arrangement is probably unlike the T cell interaction with APCs *in vivo*. Under physiological conditions, TCR present on microvilli or other membrane protrusions of T cells searches for pMHC with cognate antigens on a complex morphology of APCs. This may imply a different geometry of the TCR/pMHC (T cell/APC) interaction compared to a flat surface. Recently, diverse approaches were designed to enable pre-controlled stimulation of T cells with alternative ligand geometry. Among those, functionalised micropillars most closely mimic membrane protrusions of APCs.^2,3^ These pillars, designed to track receptor-induced forces, have a diameter of 0.5-1 μm and are more reminiscent of larger membrane protrusions such as ruffles or filopodia. Other approaches focused primarily on the spacing between stimulating nodes (pMHC clusters, e.g., <5 nm gold beads^4^) or the shape and size of stimulating ligands (e.g., using DNA origami^5^). To our knowledge, no system was developed to mimic stimulation of T cells through pMHC on microvilli of APCs. However, earlier data indicate the involvement of microvilli on dendritic cells, master APCs in humans, in the activation of T cells.^6^ Similar role of microvilli was described or predicted for other immune cells.^7–9^

Nanotechnology introduced a variety of objects with feasible sizes and functional advantages to mimic the geometry of microvilli.^10^ Nanoparticles are broadly available in a variety of compositions, sizes and surface functionalization. Metal nanoparticles, quantum dots and polymer microbeads are among the most widely used materials exhibiting either bright fluorescence or, conversely, optical transparency.^11–13^ The surface coating determines the dispersibility and stability in solvents as well as reactivity with other species. Frequently employed are inert and hydrophilic PEG chains. Desired specific reactivity is then introduced through diverse terminating groups such as carboxylic acid, thiol, azido or amino groups.^14,15^

Here we present a biomimetic platform to perform fluorescent microscopy imaging with living cells. We based the design on multilayer nano-bio platform for interactions with biological and cellular systems. Differences in the nanomaterial and biomolecule composition in turn impose challenges on the specificity and atom economy of employed transformations, which add to the mundane request for high level of control over density and distribution homogeneity of the presented activators. We were motivated to develop a practical fluorescent microscopy platform mimicking accumulation of ligands on microvilli with minimal technological requirements for a broad use. We demonstrate the functionality of the developed platform by monitoring specific T-cell activation using a cytosolic calcium-sensitive probe.

## Results and Discussion

### Design

We have designed a platform based on conventional glass coverslips for fluorescence microscopy imaging to ensure compatibility with objective working distance limitation. Glass surface functionalization is most performed by commercial triethoxysilane anchors available in a variety of functional terminations, including (3-aminopropyl)triethoxysilane (APTES) to introduce amino groups on glass. Amines on the surface allow for both irreversible and reversible ligation of small molecule linkers through alkylation and imine condensation reactions, respectively. The reversible condensation with an aldehyde grafted to the surface brings several advantages and the dual reactivity of the functionalization agent has important consequences. Firstly, during the reaction of 4-(bromomethyl)benzaldehyde with the APTES treated substrate, the irreversible nucleophilic substitution will eventually outcompete the reversible imine condensation. Secondly, both aldehyde and imine species on the surface are prone to react with amino-functionalized particles when in proximity to the surface. Thirdly, the thermodynamically governed ligation allows for chemoselective assays to confirm the nature of particle bonding to the surface as well as its ‘fixing’ by mild and chemoselective reduction to a secondary amine. Finally, optimization of desired particle density on the surface is only dependent on the particle concentration in the supernatant solution, because the particle ligation to the surface is an equilibrium of multidentate binder-substrate interaction.^16^

This primary functionalization to readily available amino functionalized nanoparticles fulfil several roles in our design: a) due to their size and curvature they mimic microvilli of a cell, which have finger-like structure with around 100 nm in diameter, b) they increase spatial separation of the presented biomolecule from the surface and thus avoid the steric clashes,^17^ and c) the amino group is suitable for biomolecule linkage onto the nanoparticle. Two types of nanoparticles, Polybead® Amino microspheres 0.10 μm polystyrene beads (PS, Polysciences) and Cd-based Amine Quantum Dots (QD, QSA580 Ocean NanoTech) were used in this study in different stages of platform development. Free surface amino groups on anchored nanoparticles were biotinylated and through tetravalent streptavidin functionalized for further attachment of desired molecules (e.g., stimulating ligands). In this work, biotinylated antibodies (anti-Hu CD3) were anchored to the particles to mimic pMHC at the tip of microvilli of APCs. Fluorescently labelled molecules were used in several stages of the development to visualize generated structures by imaging using a widefield microscope with a single-fluorophore sensitivity. An overview of the multilayer system design is presented in Scheme 1.

**Scheme 1.**
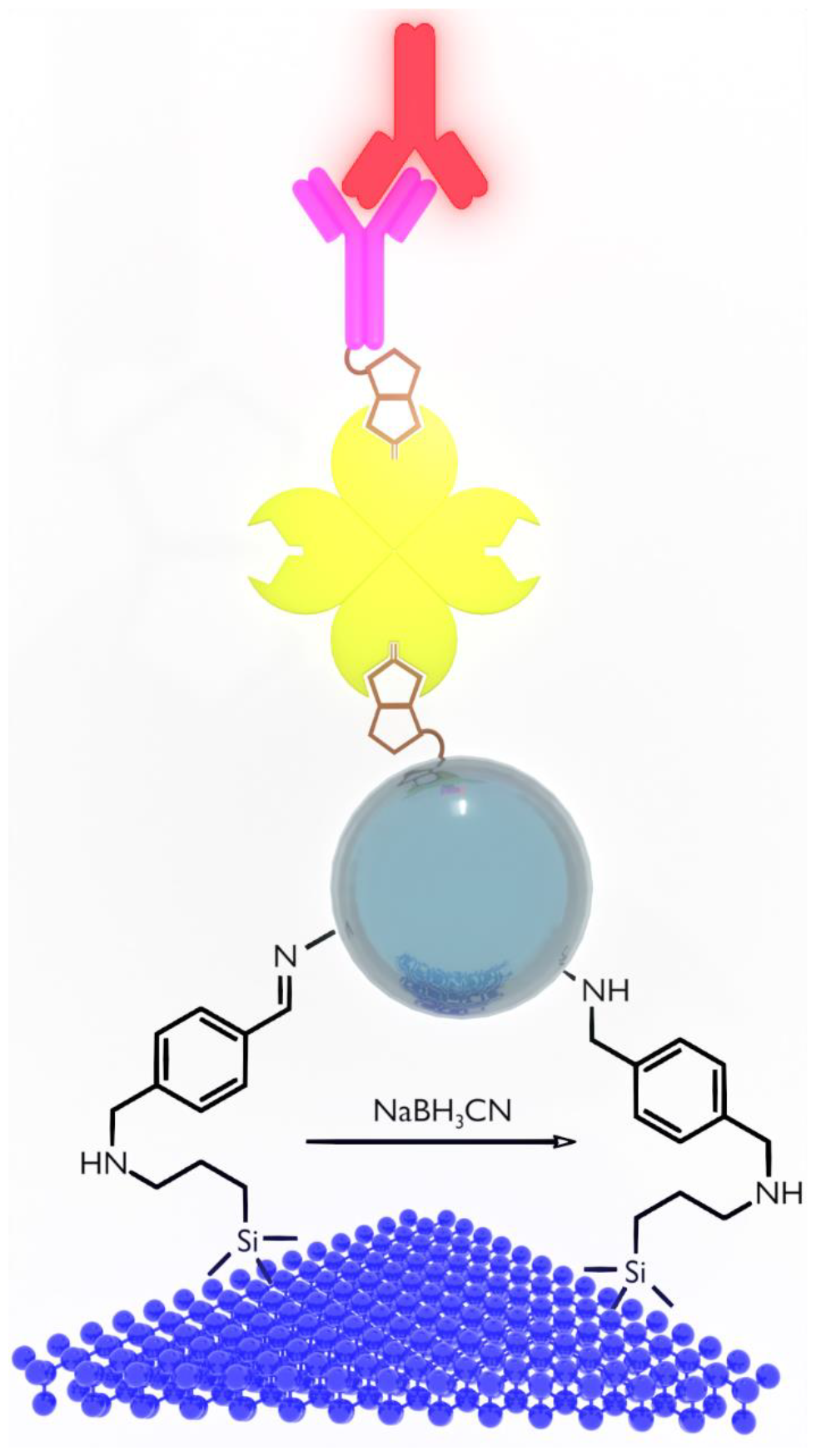
Design and development of a biomimetic nano-platform for live-cell fluorescence microscopy mimicking microvilli presentation of ligands for immune cells. Chemical surface functionalization introduces reversible imine linkage for thermodynamic binding of (nano)particles, which is fixed by mild reduction. Biotin is covalently linked to the nanoparticle and binds to tetravalent streptavidin. Free binding sites are used by biotinylated ligands (e.g., antibodies) for stimulation of imaged cells.

### Chemical surface functionalization

The coverslip surface was activated by piranha treatment to increase hydroxyl groups^18^ density for reaction with APTES.^19^ This leads to a surface with pending amino groups. To link amino-functionalized particles to such a surface, a bifunctional linker is required. We have prepared 4-(bromomethyl)benzaldehyde featuring two reactive groups. The benzylic bromide is irreversibly replaced by N-nucleophiles, such as the pending NH2 group from APTES functionalization, while the aldehyde forms imines with amine reversibly. The functionalization was monitored by measurement of contact angle (CA). ‘Piranha’-treated glass shows an immeasurably low contact angle which increases to 51 ° after APTES functionalization due to the contribution of propyl chains. After the reaction with 4-(bromomethyl)benzaldehyde, the contact angle increases further to 67 ° due to the hydrophobic nature of the benzaldehyde moiety. A summary of the basic characterization of chemically modified coverslips is shown in the Supplemental Information (SI) Figure S 1.

We attached two different types of amine functionalized particles, PS and QD, to the aldehyde-decorated glass surface of coverslips. PS are optically silent, do not interfere with the fluorescence imaging and have diameter comparable to the microvilli size (*ca* 100 nm), which is ideal for the target of this work.^20^ QD, on the other hand, are smaller (< 5 nm) and feature intense well-defined luminescence without photobleaching, which is beneficial for optimization experiments.^21^ QD were used during development of the platform for control experiments, while PS beads were the target system to mimic microvilli.

The surface bound aldehyde groups react with amine functionalities of nanoparticles to form an imine (C=N) linkage. This linkage has two major advantages. Its inherent dynamicity allows to discern non-specific sedimentation of particles on the surface from the desired covalent anchoring via chemoselective reactions. Moreover, the inherent dynamicity can be ‘frozen’ by reduction with sodium cyanoborohydride to a stable secondary amine.^22^ We have employed acetate buffer (pH 4.5) washing to remove non-bound particles. Acidic medium promotes hydrolysis of the imine linkage and protonates pendant amino groups both on the particle and the surface. This leads to Coulombic repulsion and as a result only particles covalently linked to the surface remain.

Particle density on the surface is an important parameter because crowded surface representation of antibodies would lead to a steric hindrance for cell receptor recognition. Relying on imine linkages, the ligation of particles to the surface is an equilibrium process, which can be controlled by a particle concentration in the applied dispersion. We have optimized the concentration of QD and PS dispersions to 10^15^ and 10^10^ particles/mL, respectively. After cyanoborohydride reduction and acetate buffer washing, the samples showed the desired particle density, 0.3-1 μm^-2^ on the surface, as characterised by AFM, and for QDs also long-term fluorescence, thus demonstrating stability of the particle anchoring (Figure 1).^23^ This density represents 10-80 particles per the area, which forms a contact site between a lymphocyte and the glass surface. It is within the range of expected microvilli contacts between a lymphocyte and a target cell.^6^

**Figure 1.**
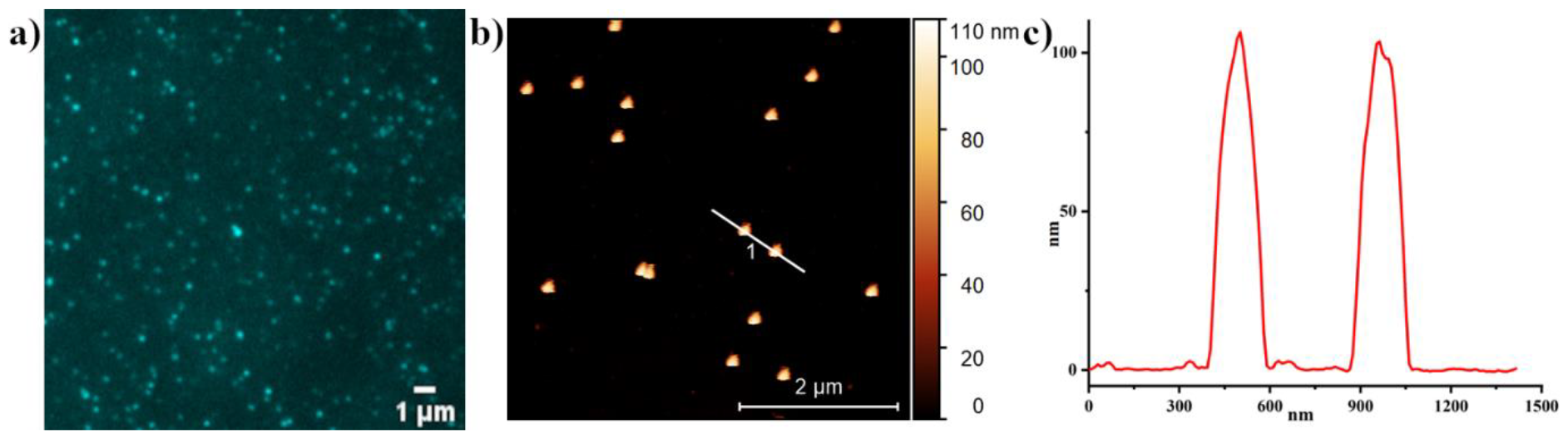
a) Fluorescence image of a coverslip surface with anchored QD, excitation wavelength 405 nm, cut-off filter 525 nm. b) AFM image of a coverslip surface with PS beads showing on average 0.7 PS bead per μm^2^. c) Height AFM profile of marked line in b showing particle height/size of 100 nm, i.e. similar to the microvilli diameter.

### Biomolecule immobilization

Attachment of signalling biomolecules to the particle surface was pursued via strong, specific biotin-avidin binding. First, the particle surface was biotinylated with 15 mM in aqueous solution of sulfo-NHS-biotin (30 minutes at r.t.), which reacts with pending NH2 group forming a stable amide bond.^24^ To confirm successful biotinylation, we have incubated both the QD and PS-decorated surface with 20 ng/ml Alexa Fluor™ 647-labelled streptavidin. In both cases, characteristic fluorescence of the dye was recorded in the micrograph (Figure 2). In contrast to QDs, the fluorescence signal of streptavidin was dramatically reduced upon photobleaching (SI Figure S 3). Inherent QD signal, on the other hand, did not undergo photobleaching and their signal remains stable upon bleaching energies applied to the sample. The results are summarized in Figure S 3. Using the fluorescently labelled streptavidin, we have optimized its concentration to 20 ng/ml in aqueous phosphate-buffered saline (PBS) with 0.5% bovine serum albumin (BSA, see SI section 2 for details).

**Figure 2.**
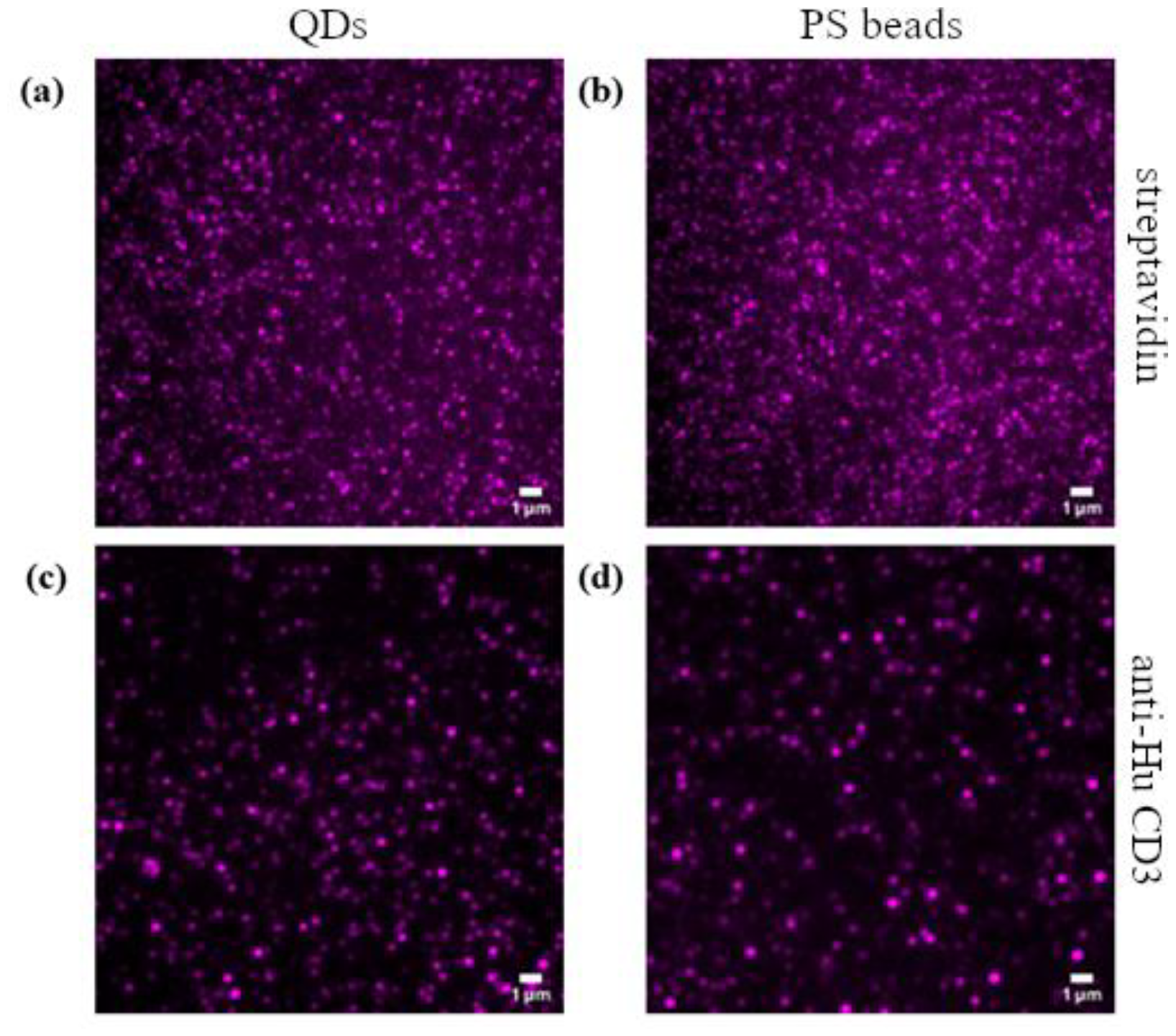
Visualization of QD (a and c) and PS beads (b and d) on functionalized nano-bio platform by fluorescence imaging: (a) & (b) represents binding of Alexa Fluor™ 647-labelled streptavidin (see Scheme 1 for details); (c) & (d) represents binding of anti-Hu CD3ε biotin (mouse IgG) via non-labelled streptavidin and visualized by goat anti-mouse IgG Alexa Fluor 647-labelled secondary antibody.

To test the final step of the nano-bio platform functionalisation, we have incubated the system with biotinylated anti-human CD3ε (MEM-57) antibody, which binds to the remaining sites of tetravalent streptavidin. The MEM-57 antibody binding was confirmed and visualized by complexation of this primary antibody with Alexa Fluor™ 647-labelled goat anti-mouse IgG antibody (secondary antibody). Fluorescence images of the system after staining with the secondary antibody demonstrate successful formation of the fully functionalised nano-bio platform for stimulation of T cells (Figure 2). Photobleaching experiments were performed to confirm the presence of the organic dye – Alexa Fluor 647 (SI Figure S 4).

### Live-cell imaging of T-cell activation on functionalised nano-bio platform

To demonstrate successful development of the platform for T-cell activation, we monitored calcium mobilisation in the immortalised model of T cells (Jurkat E6 cell line, ATCC TIB-152^25^) expressing GCaMP6f genetically encoded calcium sensor.^26^ Jurkat cells forming contact with nanobeads functionalised with activating anti-human CD3ε antibody exhibited transient, but strong increase in the intracellular calcium (SI suppl_vid_01, suppl_vid_02, suppl_vid_03). Almost 60 % of cells on the nano-bio platform with anti-human CD3ε antibody underwent activation whereas only approximately 20 % of cells on the surface generated by omitting the last functionalisation step with the antibody (negative control) exhibited a weak calcium mobilisation (Figure 3). This was probably caused by the mechanical stress upon T cell contact with the coverslip in the unmodified areas. Detailed image analysis demonstrates active engagement of membrane protrusions on the surface of Jurkat cells with the activating surface (Figure 3c). No such preferred morphology was detected in cells on non-stimulating surface (Figure 3b, SI suppl_vid_04, suppl_vid_05, suppl_vid_06).

**Figure 3.**
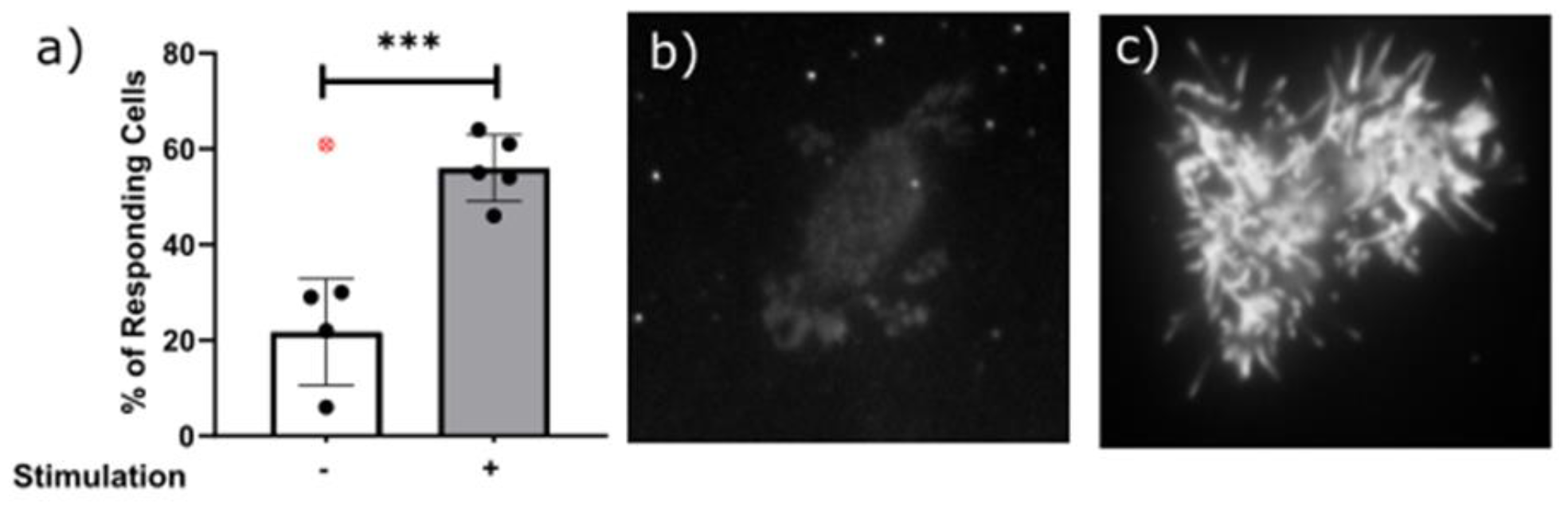
**a)** Calcium response of Jurkat cells upon contact with the nano-bio platform containing QD lacking antibodies (-) (n = 370) or QD with anti-human CD3ε antibodies (MEM-57) (+) (n = 218). The magnitude of the calcium was calculated using the software CalQTrace.^27,28^ Relative calcium response was calculated by an AI-based decision algorithm to classify individual cells as triggering (activated) or non-triggering (resting) upon contact with the modified coverslip and during 15 min of the measurement at 37 °C. Statistical difference was assessed using unpaired T-test (GraphPad Prism 9); ***P < 0.001. Dots in the graph represent independent measurements. Red crossed dot represents a value of an outlier. b) and c) Representative images of a Jurkat cell contact with non-stimulating (b) and stimulating (c) nano-bio platform. Cells were transfected with genetically encoded calcium sensor GCaMP6fu and imaged using total internal reflection fluorescence (TIRF) microscopy. Full time-lapse series of the live-cell imaging of Jurkat cell activation on nano-bio platform are in SI suppl_vid movies.

## Conclusions

The antibody-functionalized platform with T-cell stimulating capacity has been fabricated through simple and straightforward protocols and a design schematically represented in Scheme 1. It combines the characteristics of ‘off-the shelf’ availability, design versatility and multiple interactions to achieve highly efficient T-cell activation. The approaches to modulate antigen-specific T cell responses presented here are also meant to be used as a guideline to develop various immune cell stimulating or modulating that imitate diverse conditions of the human immune system. In this work, we demonstrated principal function of our system by focusing on a nanopatterned T-cell activating platform on an optical surface, which allows multiplexed imaging with selected ligands. Further development is required to combine stimulating particles with integral negative control or dual-target platform.

## Supporting information

suppl_vid_01

suppl_vid_02

suppl_vid_03

suppl_vid_04

suppl_vid_05

suppl_vid_06

Summplemental information

## Acknowledgements

The work was supported by the Czech Science Foundation Grant No. 22-11299M ‘Reaction networks at phase interfaces for dynamic self-assembly’. The team was also supported by the Experientia Foundation 2021 start-up grant. We acknowledge the support of the Dagmar Prochazkova Foundation of the University of Chemistry and Technology Prague. The work of J.A.G.N., D.A., T.C., T.R.B.R. and M.C. was also supported by the Czech Science Foundation grant No. 19-07043S.

