## Supplementary material for "Cell surface morphology mimicking nano-bio platform for immune cell stimulation": Summplemental information

### Supplementary Information file: Cell surface morphology mimicking nano-bio platform for immune cell stimulation

#### 1 Instrumentation

##### 1.1 Contact angle measurement

Contact angle goniometer (Ossila Ltd, UK) was used to measure the contact angle in static mode at room temperature. A drop of deionized water was gently placed onto the surface (averages of n = 3 determinations) and analysed by Ossila contact angle software.

##### 1.2 Atomic Force Microscopy

The topological images of the PS beads functionalized surface were achieved by A Nanosurf NaioAFM (Nanosurf, Switzerland), operated in dynamic AFM mode at room temperature. The imaging was performed using Dyn 190 Al tips in the measurement area of 10 μm ×10 μm and were zoomed in for getting more precise images. The images were collected at a scanning rate of 1 Hz with resolution of 512×512 points and were processed using Gwyddion software.[29]

##### 1.3 Fluorescence Microscopy

Fluorescence microscopy was performed on a home-built single-fluorophore sensitivity wide-field microscope with TIRF modality based on the IX71 body (Olympus). Fluorescence of QD QSA580 (Ocean NanoTech) was excited by 150 mW 405 nm laser (Cube, Coherent) and of Alexa Fluor™ 647 Conjugated streptavidin as well as IgG Alexa Fluor 647 antibody by 170 mW 638 nm laser (Lasos), respectively. All laser lines were focused to the back focal aperture of a 100x oil immersion TIRF objective (UApo N 100x, 1.49 NA, Olympus). TIRF illumination was achieved using a manual large-area translation platform (Thorlabs). Excitation and emission were separated through a filter-cube (Olympus) equipped with quadband beamsplitter (Chroma) and quadband emission filter (Chroma). Emission was additionally filtered out with 525/50 filter (Semrock) for Qdots, 605/70 or 655 LP (Chroma) for Alexa Fluor 647 positioned in front of EMCCD camera (iXon DU-897U, Andor). EM gain of the camera was set to 300, 100 images were acquired with the exposure time of 50 ms and summed in post-processing. Two acousto-optic tuneable filters (AOTFnC-400.650-TN, AA Optoelectronics) provided fast switching and synchronization of lasers with a camera. The whole system was controlled by µManager software.

#### 2 Chemical Reagents & Solutions

Ammonium hydroxide ($NH_4OH$), Ethanol (EtOH, UV grade), 3-aminopropyltriethoxysilane (APTES), Acetonitrile (ACN, UV grade), sodium cyanoborohydride ($NaBH_3CN$) (Penta Chemicals, Czech Republic), hydrogen Peroxide ($H_2O_2$) (Lach:ner, Czech Republic, Polybead® amino microspheres, 0.112 µm (Polysciences, Inc., Germany), Qdot™ 525 ITK™ Amino (PEG) quantum dots, sulfo-NHS-biotin, streptavidin, Dylight™Alexa Fluor 647 streptavidin, Alexa Fluor 647 goat anti-mouse IgG2A (Y2a) (Thermo Fisher Scientific, USA), anti-Hu CD3 biotin (MEM-57, Exbio, Czech Republic), were purchased and used as received without further purification.

Ultrapure water was obtained in the laboratory using a Milli-Q water purification system. 1 mM solution of (**1**) was prepared in acetonitrile and kept at room temperature. 1 µl in 5 ml was optimized as the concentration of nanoparticle solution and stored at 4 °C. Just prior to use, different stock solutions of sodium cyanoborohydride (13 mg) and biotin (2 mg) were prepared in

300 µl and 1000 µl of water, respectively. 0.5% BSA in PBS was used to prepare streptavidin and antibody solutions and to prevent non-specific interactions of the molecules with the surface. Sodium acetate buffer (0.2 M, pH 4.5) was prepared by dissolving sodium acetate trihydrate in distilled water; the final pH was adjusted by acetic acid (0.2 M). Phosphate-buffered saline, 10×PBS contains 80 g sodium chloride, 2 g potassium chloride, 14.4 g disodium hydrogen phosphate and 2.4 g potassium dihydrogen phosphate in 1 litre of Milli-Q water. The mixture was sterilized by autoclaving, stored at room temperature and diluted 10× prior to use to get 1×PBS used throughout the study. Bovine serum albumin, BSA, solutions were prepared by directly dissolving BSA in 1×PBS. It was kept at 4 °C. All buffers were filtered through 0.22 µm nylon membrane filter.

### 3 Methods

#### 3.1 Surface functionalization

**Cleaning**: In terms of high-resolution imaging, biocompatibility as well as degradability, optical coverslips (25 mm) are selected as an ideal platform for fabricating the multilayer system. The cleaning methods include the sonication of coverslips in ultrapure water, acetone, and isopropyl alcohol, consecutively for 10 minutes. After washing and heating at 80 °C in diluted basic Piranha solution (5:1:1 v/v mixture of water, 23% aqueous ammonia and 30% aqueous hydrogen peroxide, respectively) for 20 minutes, the coverslips are rinsed with distilled water and dried.

**Amino silanization**: The cleaned coverslips were placed in a petri dish with 3% APTES in ethanol at room temperature. After 20 minutes, the surface was thoroughly rinsed with ethanol and dried completely.

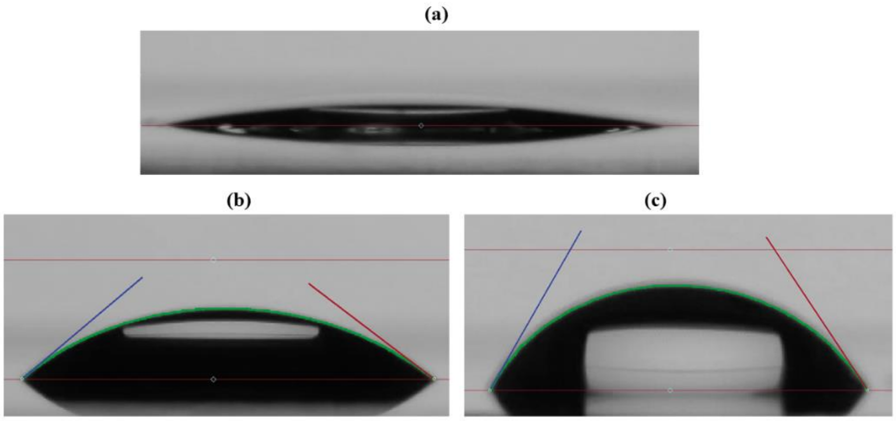

*Figure S 1 water contact angle on glass surface after treatment with (a) Piranha solution (b) APTES (c) Aldehyde.*

**Treatment with (1):** The APTES modified substrate was kept in solution of 4-(bromomethyl)benzaldehyde for 45 minutes, followed by washing with distilled water and drying.

**Functionalization with PS beads and Qdots**: The nanoparticle solution (200 µl) followed by $NaBH_3CN$ (300 µl) were separately pipetted on the aldehyde functionalized surface at room temperature for 45 minutes. After washing and drying, the coverslip was immersed in acetate buffer for 30 minutes. Then the surface was subsequently washed with distilled water and dried.

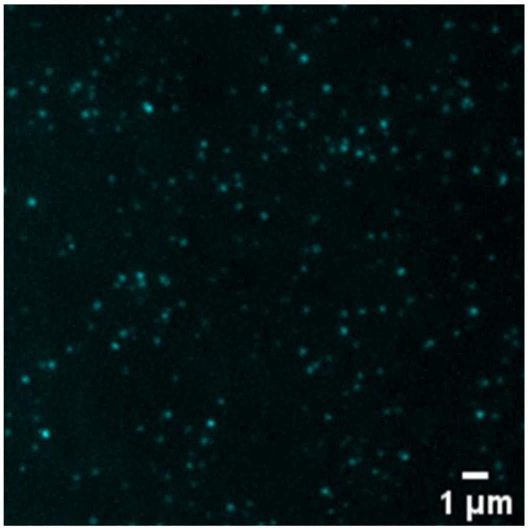

*Figure S 2 FlM image on coverslip surface with Qdots after washing (45 minutes); QDs 1 µl in 5 ml of $H_2O$, exposure time 50 ms, EM Gain 300, 100 frames summed up, excitation 405 nm, longpass filter cut-off 500 nm, monochromatic image colorized in Fiji/ImageJ.*

**Bioconjugation with biotin-streptavidin**: Biotinylation was achieved by adding 300 µl of the biotin solution onto the nanoparticle functionalized surface for 30 minutes at room temperature. Afterwards, the surface was washed with water, methanol, and water. Finally, the coverslips were incubated in 1% BSA-1×PBS at 37 °C for 1 h to block the non-specific binding site. After washing with water, the nanosurface was incubated with 300 µl of unlabelled streptavidin solution (10 ng/ml) at room temperature for 30 minutes, followed by washing with 300 µl of 0.5% BSA in PBS (5×3 minutes).

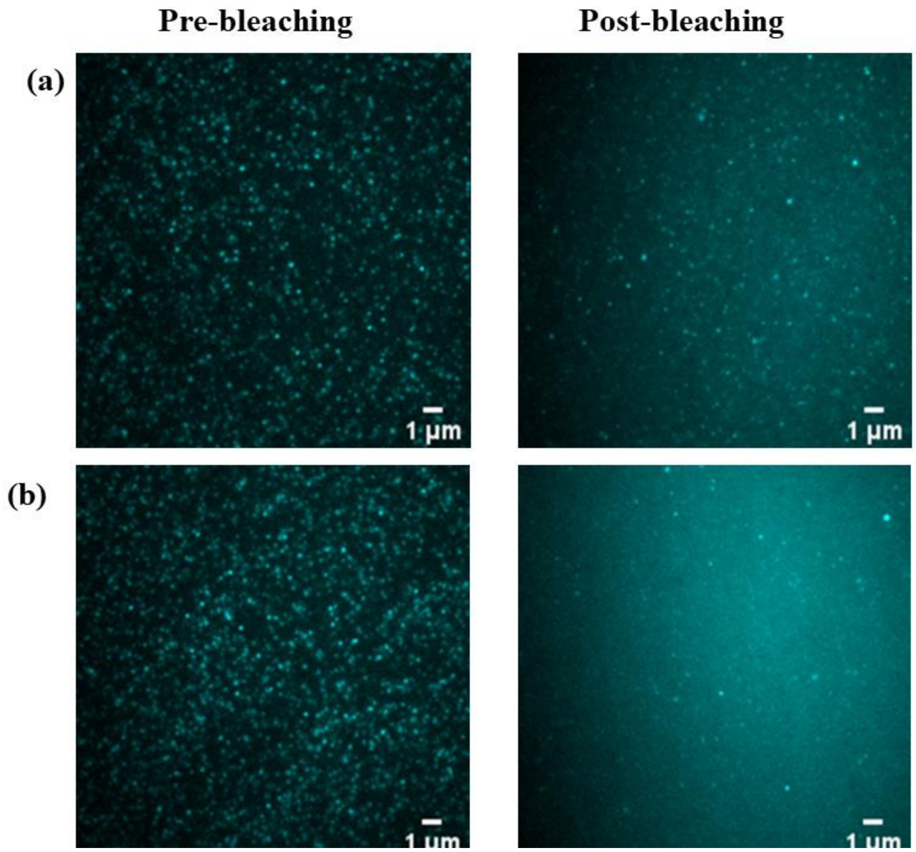

*Figure S 3 Fluorescence images of Alexa Fluor™ 647-streptavidin bound to immobilised and functionalised: (a) QD (b) PS beads on glass coverslips (see Scheme 1 for details), excitation 638 nm, longpass filter cut-off 648 nm, monochromatic image colorized in Fiji/ImageJ.*

**Antibody immobilization**: The antibody immobilization was performed by exposing the functionalized surface to 300 µl of antibody solution on parafilm for 30 minutes at room temperature. After incubation, the surface was thoroughly washed with 0.5% BSA in PBS (5×3 minutes) and another blocking with 5% BSA in PBS at 37 °C for 30 minutes.

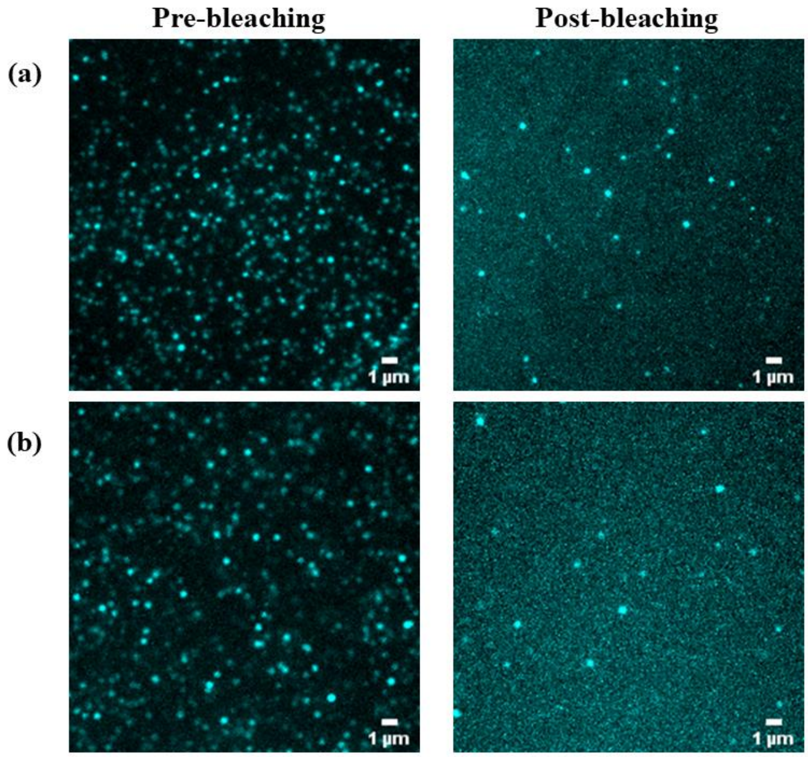

*Figure S 4 FlM images of the microvilli mimicking platform after binding of IgG Alexa 647 secondary antibody. (a) QD (b) PS beads, excitation 638 nm, longpass filter cut-off 648 nm, monochromatic image colorized in Fiji/ImageJ.*

##### Cell culture and transfection.

Jurkat cells (clone E6-1; ATCC) were grown in RPMI-1640 medium (Fischer Scientific) supplemented with 10% of foetal bovine serum (Fischer Scientific), non-essential amino acids (Fischer Scientific) and 10mM HEPES under controlled humidity in the incubator at 37 °C and 5 % $CO_2$. The cell line was regularly tested for morphology and mycoplasma infection.

For the expression of GCaMP6fu and GCaMP6fast genetically encoded calcium probes, Jurkat cells were transiently transfected with DNA plasmids using the Neon® transfection system (Thermo Fisher Scientific). The transfection was performed using 1 µg of the DNA plasmid and 300,000 cells in 0.5 ml of pre-warmed supplemented RPMI-1640 culture medium. The instrument settings were 3 pulses, each 1500 V for 10 ms. The cells were incubated for 16-24h before their use.

**Large-scale calcium measurement.**

The functionalized coverslips were mounted in a ChamLide holder (Live Cell Instruments) with a microfluidics system and kept in 500 ul of PBS-BSA 5%, attached in a live-cell environmental chamber (Uno, Okolab S.r.l.) at 37 °C for the entire experiment.

A day after transfection, 600.000 cells were centrifuged for 3 min at 0.3 rcf and resuspended in 1 ml of pre-warmed colour-free RPMI-1640 supplemented with 10% of foetal bovine serum (Fischer Scientific), non-essential amino acids (Fischer Scientific), 1 mM $CaCl_2$ and 1 mM $MgCl_2$. Immediately after resuspension, the cells were transferred to the coverslip using the microfluidics system, with the image acquisition running shortly before.

Live-cell calcium influx measurements were performed on the IX73 frame (Olympus) equipped with ×20 objective (UPLXAPO20X, Olympus), LED epifluorescence illumination system (CoolLED pE-4000, CoolLED) and sCMOS camera (Zyla 4.2P-CL10, Andor). Coated coverslips were mounted into the ChamLide holder with a microfluidics system attached to a live-cell environmental chamber (Uno, Okolab S.r.l.) set to 37 °C. The acquisition time for one frame was 250 ms with illumination using 470 nm laser set to 10%. Cells were imaged for 15 min (3600 frames) after injection into the imaging chamber. The acquisition was controlled by μManager software (ver. 1.4.24).[30]

Data analysis. Obtained data were converted to 8-bit mode and processed to have a minimum intensity of 20-35 using Fiji/ImageJ software (version 1.53f51),[31] not affecting the brightness of the cells and evenly reducing the background variations. The data was saved and uploaded to the nobel AI-algorithm CalQTrace (unpublished) based on the previously reported algorithm CalQuo2[28] to calculate the calcium response of single cells. The algorithm was specified to detect cells with a 24-26 pixels diameter, with a brightness slightly higher than the minimal background, single cells were tracked for at least 1500 frames (6 min, 15 sec), and the algorithm classified the cells depending on the variations of the normalized intensity they presented. Five independent experiments were performed. Statistical differences were evaluated using an unpaired T-test; $^*P < 0.05$ (GraphPad Prism 9).

**Single-cell calcium measurement.**

The coverslips were mounted to the ChamLide imaging chamber, and the cells were transferred to the coverslip as previously described for the large-scale calcium measurement. Cells transfected with genetically-encoded calcium probe GCaMP6fast were imaged using a home-built super-resolution wide-field TIRF microscope described in section 1.3. The imaging was performed in the TIRF mode using the 488 nm laser with a 5% power intensity; the emission was filtered using 525/50 filter (Semrock) positioned in front of the EMCCD camera (iXon DU-897U, Andor). EM gain of the camera was set to 100 with an exposure time of 100 ms; individual cells were imaged for at least 15 min. The calcium influx was analyzed by the mean fluorescence intensity (MFI) changes of the cells landing on the coverslips as previously described.[21]

*Table S 1 Descriptive statistics of large-scale measurement calcium influx*

| | Stimulus (-) (n=370) | Stimulus (+) (n=218) |
|---|---|---|
| Number of values | 5 | 5 |
| Minimum | 6.000 | 46.00 |
| 25% Percentile | 14.00 | 50.00 |
| Median | 29.00 | 55.00 |
| 75% Percentile | 45.50 | 62.50 |
| Maximum | 61.00 | 64.00 |
| Range | 55.00 | 18.00 |
| 10% Percentile | 6.000 | 46.00 |
| 90% Percentile | 61.00 | 64.00 |
| 95% CI of median | | |
| Actual confidence level | 93.75% | 93.75% |
| Lower confidence limit | 6.000 | 46.00 |
| Upper confidence limit | 61.00 | 64.00 |
| Mean | 29.60 | 56.00 |
| Std. Deviation | 20.01 | 6.964 |
| Std. Error of Mean | 8.948 | 3.114 |

| | | |
|---|---|---|
| Lower 95% CI of mean | 4.757 | 47.35 |
| Upper 95% CI of mean | 54.44 | 64.65 |
| Coefficient of variation | 67.59% | 12.44% |

*Table S 2 Outlier test of large-scale measurement calcium influx (ROUT Q = 10%)*

| | - (n= 370) | + (n=218) |
|---|---|---|
| Method | | |
| ROUT (Q = 10%) | | |
| Number of points | | |
| # Y values analyzed | 5 | 5 |
| Outliers | 0 | 0 |

*Table S 3 Summary table of results of unpaired t-test of large-scale measurement calcium influx.*

| | |
|---|---|
| Table Analyzed | Less one Percentage of responding cells |
| Column B | + (n=218) |
| vs. | vs. |
| Column A | - (n= 370) |
| Unpaired t test | |
| P value | 0.0007 |
| P value summary | *** |
| Significantly different (P < 0.05)? | Yes |
| One- or two-tailed P value? | Two-tailed |
| t, df | t=5.694, df=7 |

| | |
|---|---|
| How big is the difference? | |
| Mean of column A | 21.75 |
| Mean of column B | 56 |
| Difference between means (B - A) ± SEM | 34.25 ± 6.015 |
| 95% confidence interval | 20.03 to 48.47 |
| R squared (eta squared) | 0.8225 |
| F test to compare variances | |
| F, DFn, Dfd | 2.534, 3, 4 |
| P value | 0.3906 |
| P value summary | ns |
| Significantly different (P < 0.05)? | No |

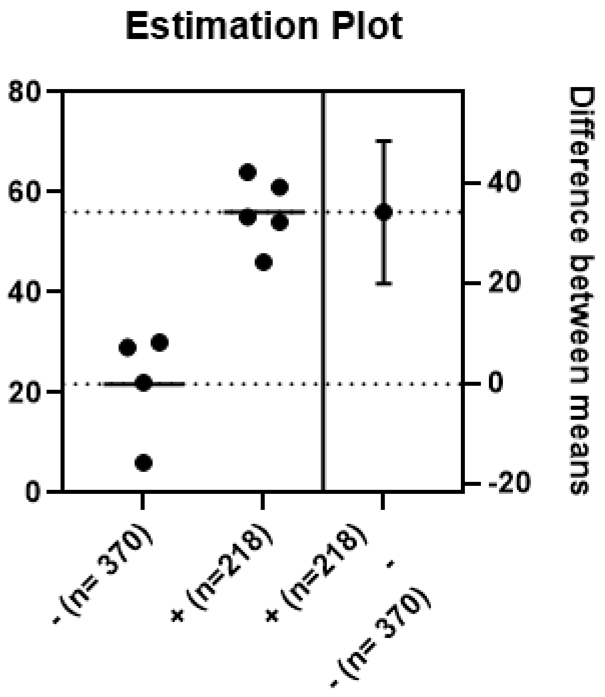

*Figure S 5 Estimation plot unpaired t-test of large-scale measurement calcium influx.*

*Table S 4 Normality test of large-scale measurement calcium influx.*

| | - (n= 370) | + (n=218) |
|---|---|---|
| Compare normal and lognormal | | |
| Probability normal (Gaussian) | 71.74% | 52.56% |
| Probability lognormal | 28.26% | 47.44% |
| Likelihood ratio (LR) | 2.538 | 1.108 |
| 1/LR | 0.394 | 0.9026 |
| Which distribution is more likely? | Normal | Normal |
| Shapiro-Wilk test | | |
| W | 0.8456 | 0.9609 |
| P value | 0.2124 | 0.8139 |
| Passed normality test (alpha=0.05)? | Yes | Yes |
| P value summary | ns | ns |
| Kolmogorov-Smirnov test | | |
| KS distance | N too small | 0.187 |
| P value | | >0.1000 |
| Passed normality test (alpha=0.05)? | | Yes |
| P value summary | | ns |
| Shapiro-Wilk test | | |
| W | 0.7656 | 0.9491 |
| P value | 0.0535 | 0.7305 |
| Passed lognormality test (alpha=0.05)? | Yes | Yes |
| P value summary | ns | ns |
| Number of values | 4 | 5 |
| Impossible values in lognormal distributions | | |
| Number of zeroes | 0 | 0 |
| Number of negative values | 0 | 0 |
